## Supplementary material for "The landscape of antibiotic resistance genes in the nasopharynx of preterm infants: Prolonged signature of hospitalization and effects by antibiotics": All the supplementary Figures

Supplementary Fig. 1

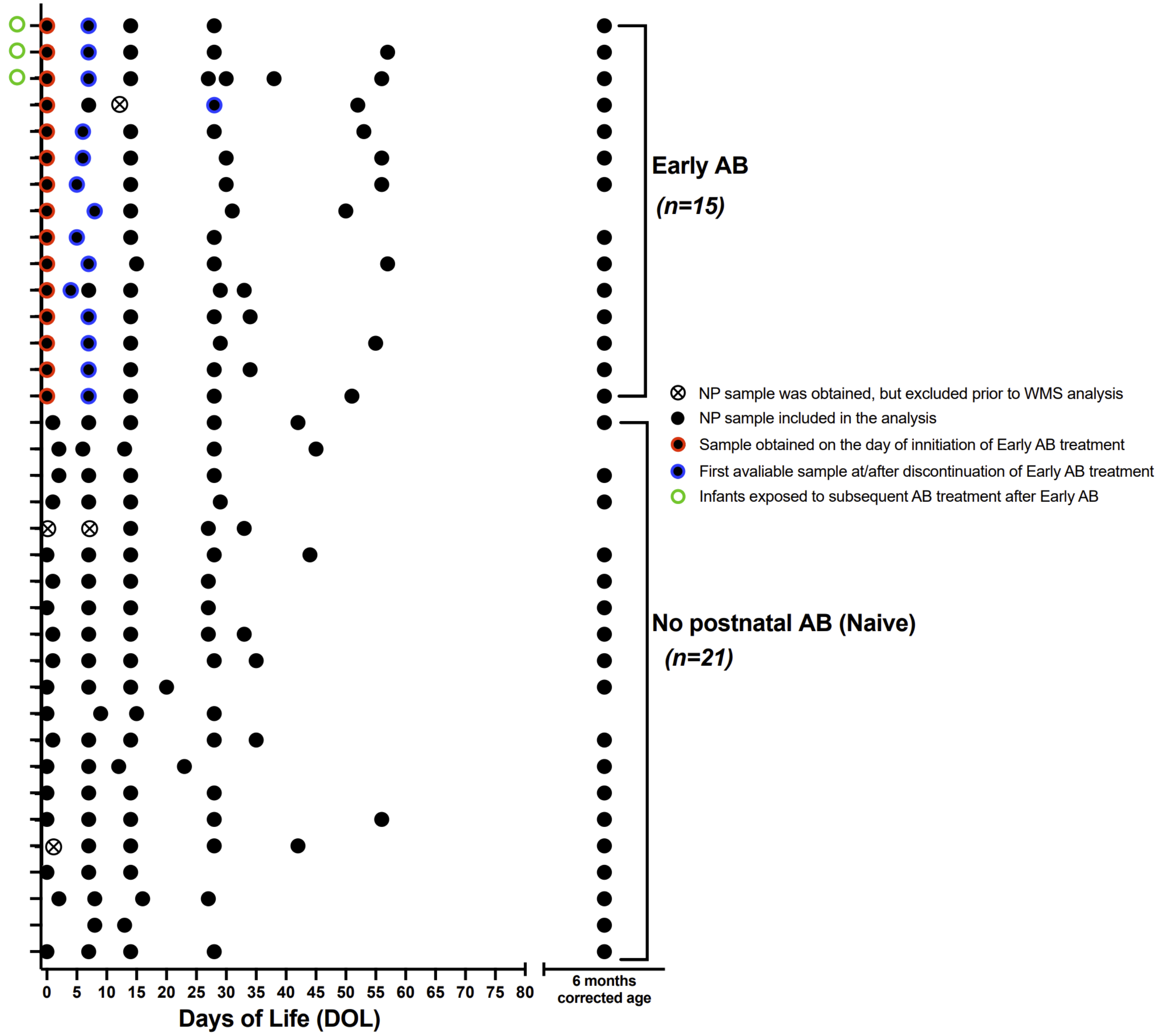

**Supplementary Fig. 2**

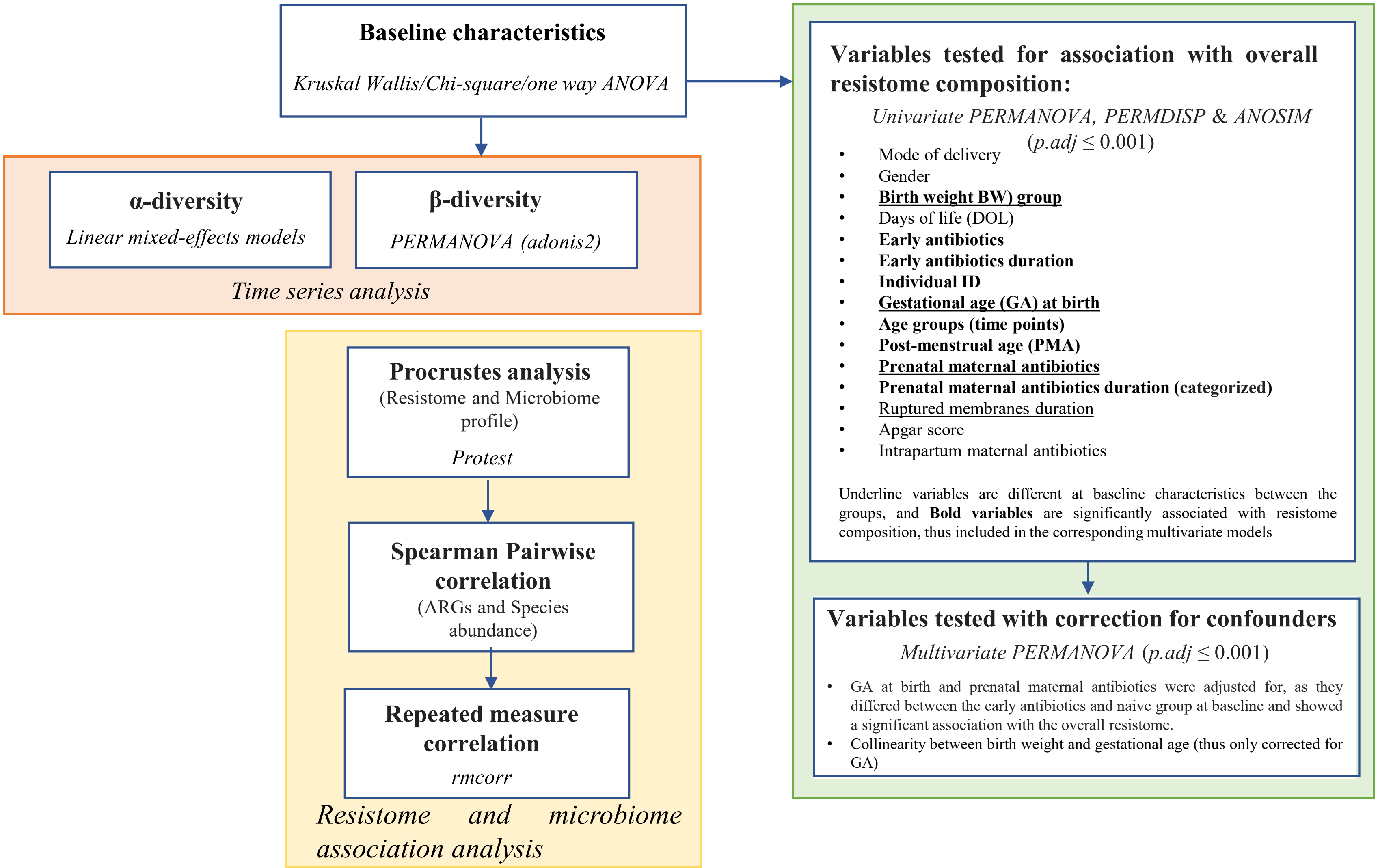

### Supplementary Fig. 3

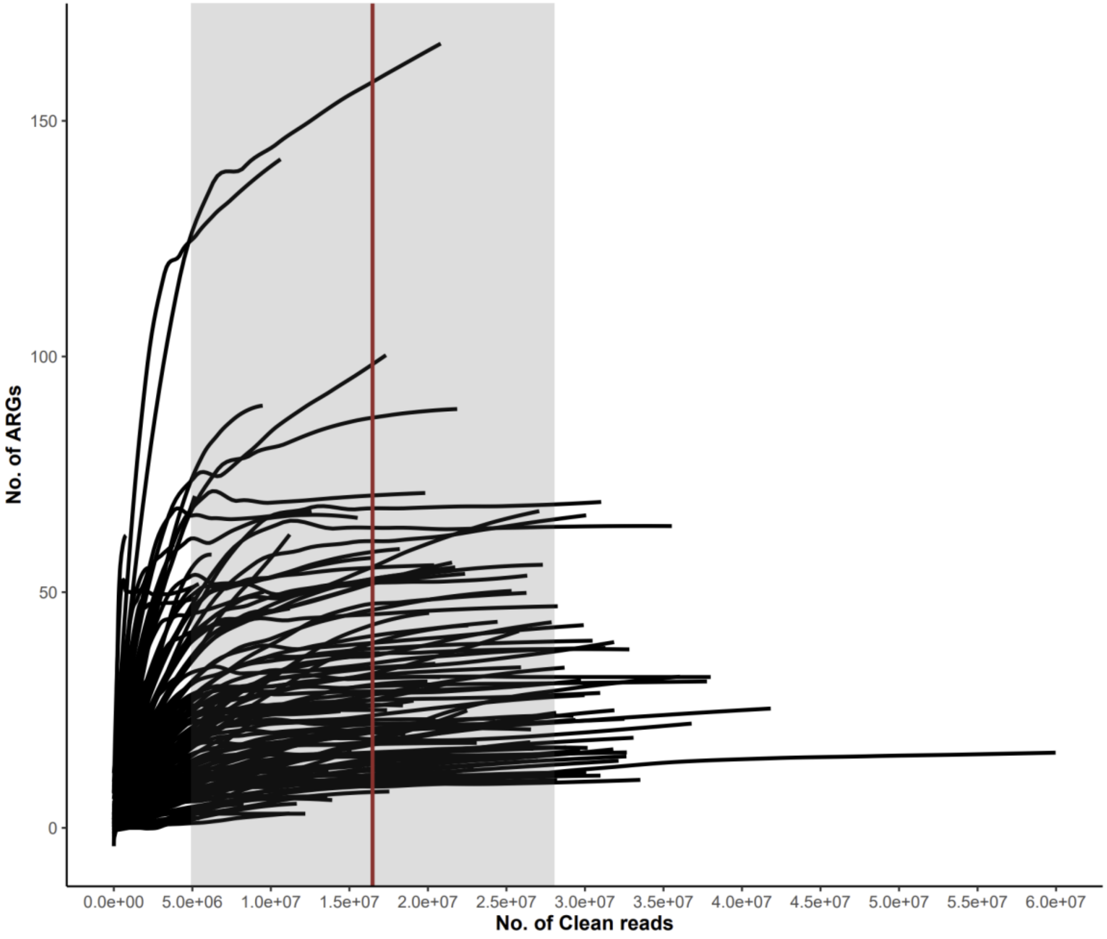

**Supplementary Fig. 4**

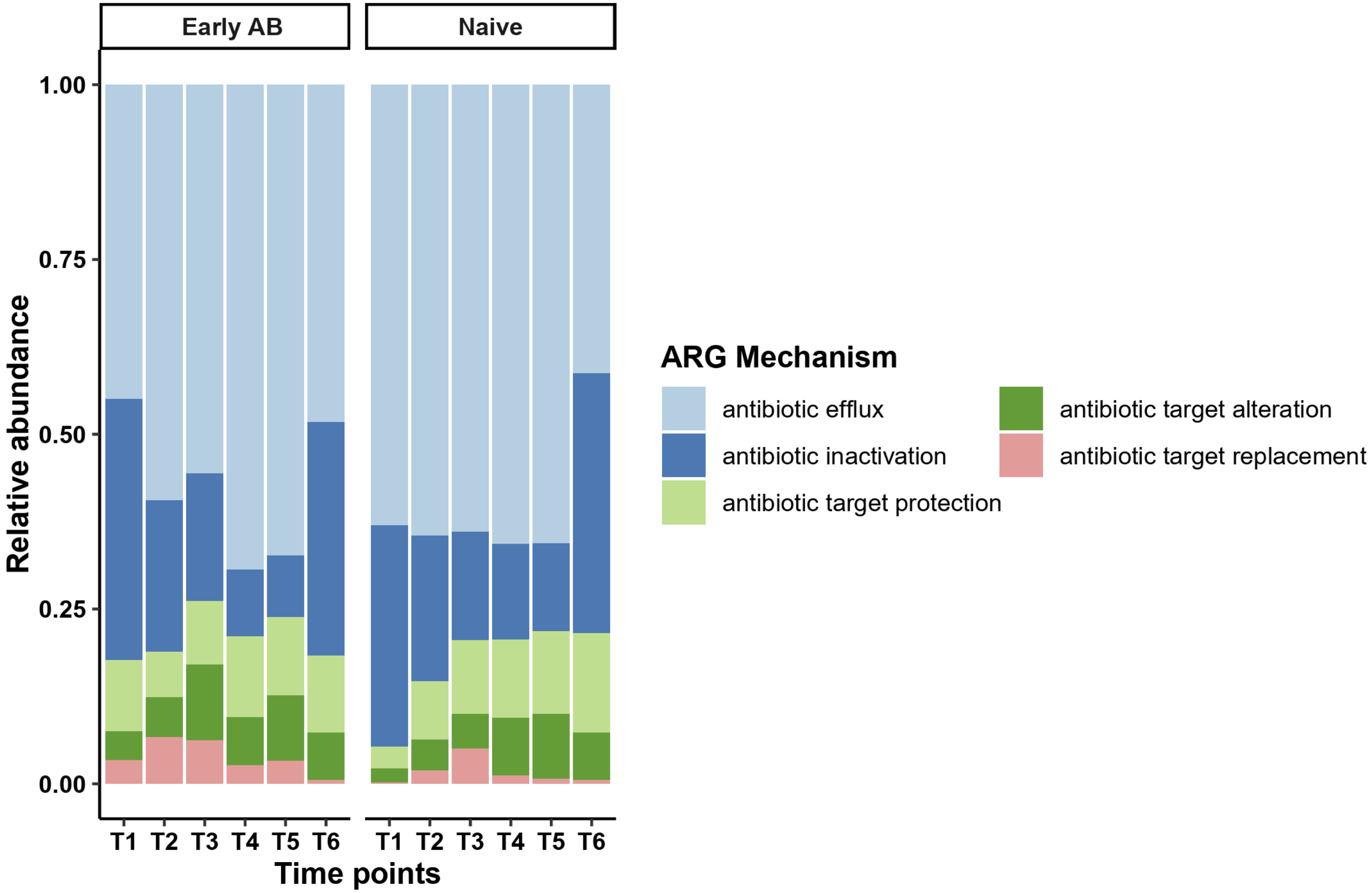

**Supplementary Fig. 5**

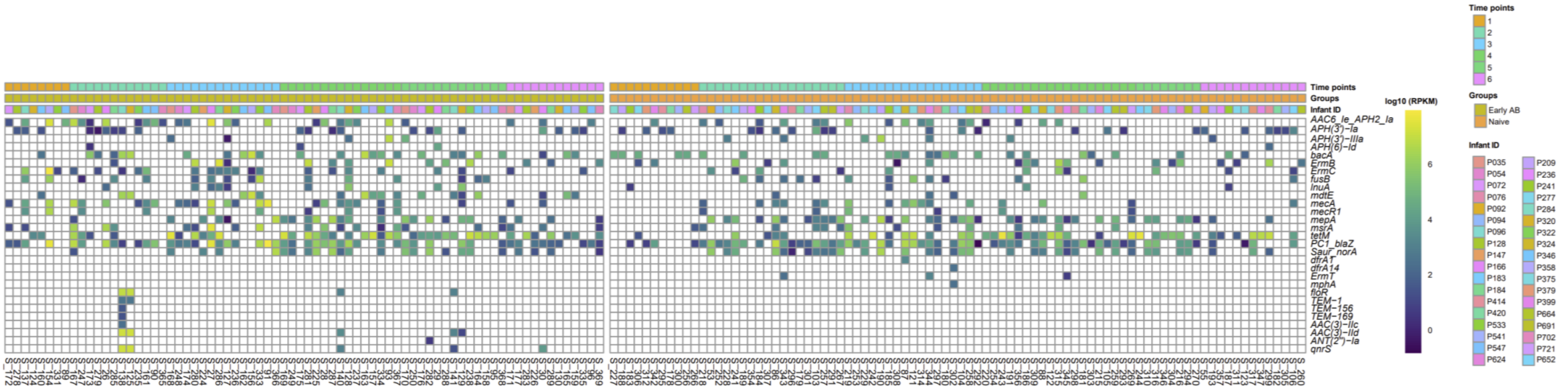

**Supplementary Fig. 6**

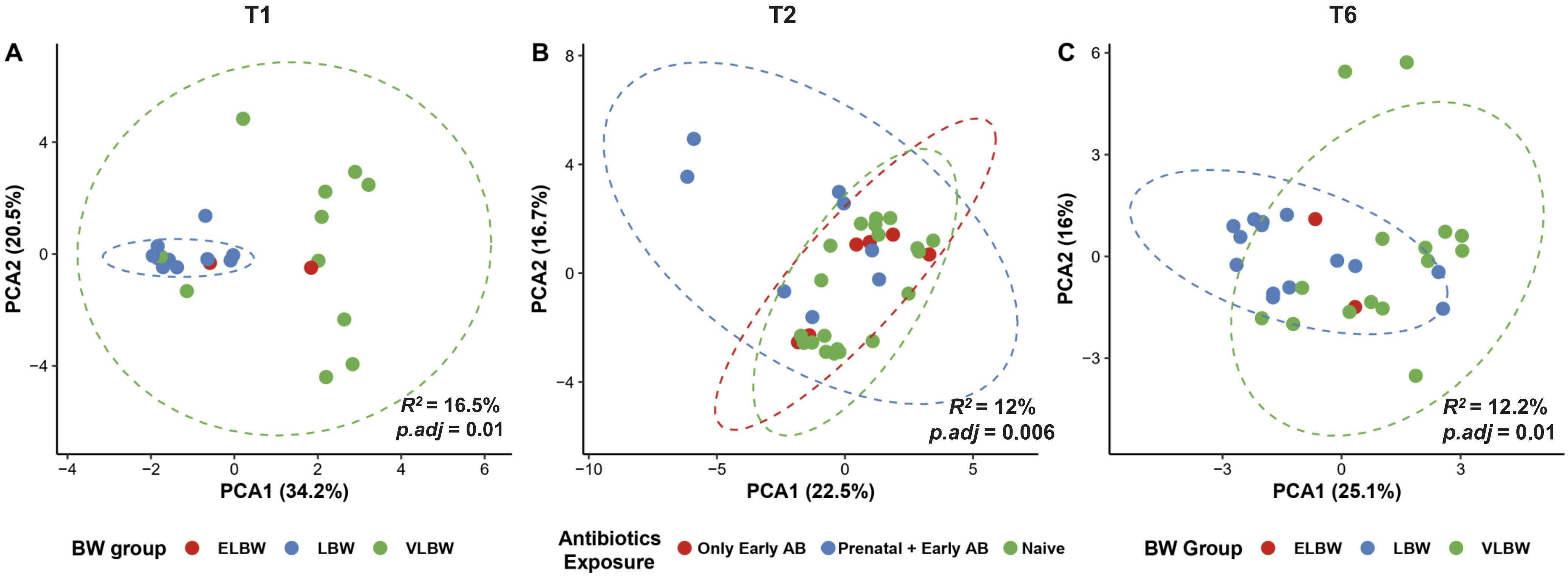


(Prenatal + Early AB Vs naive): $R^2$ = 10.1%, *p.adj* = 0.009

**Supplementary Fig. 7**

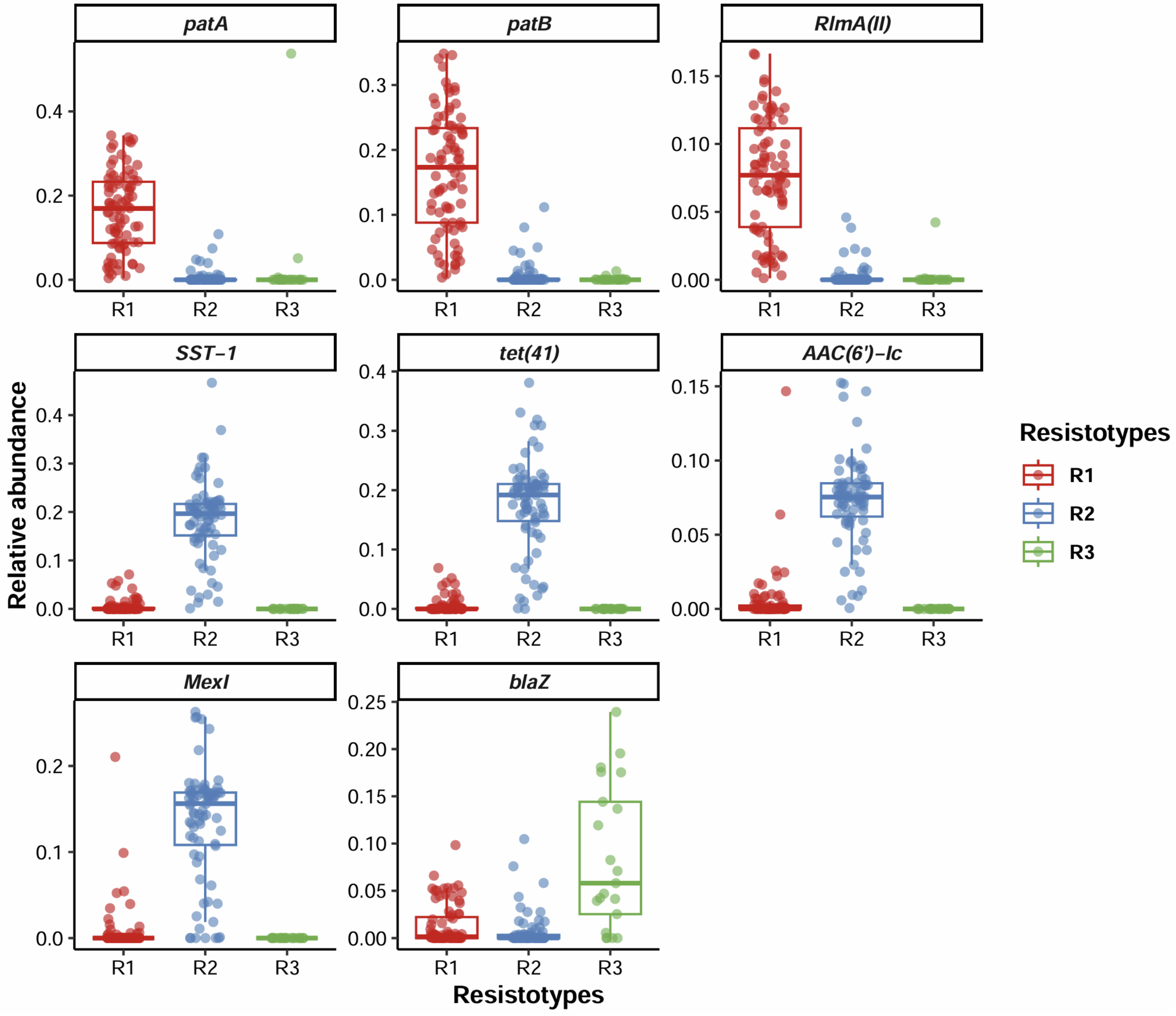

Supplementary Fig. 8

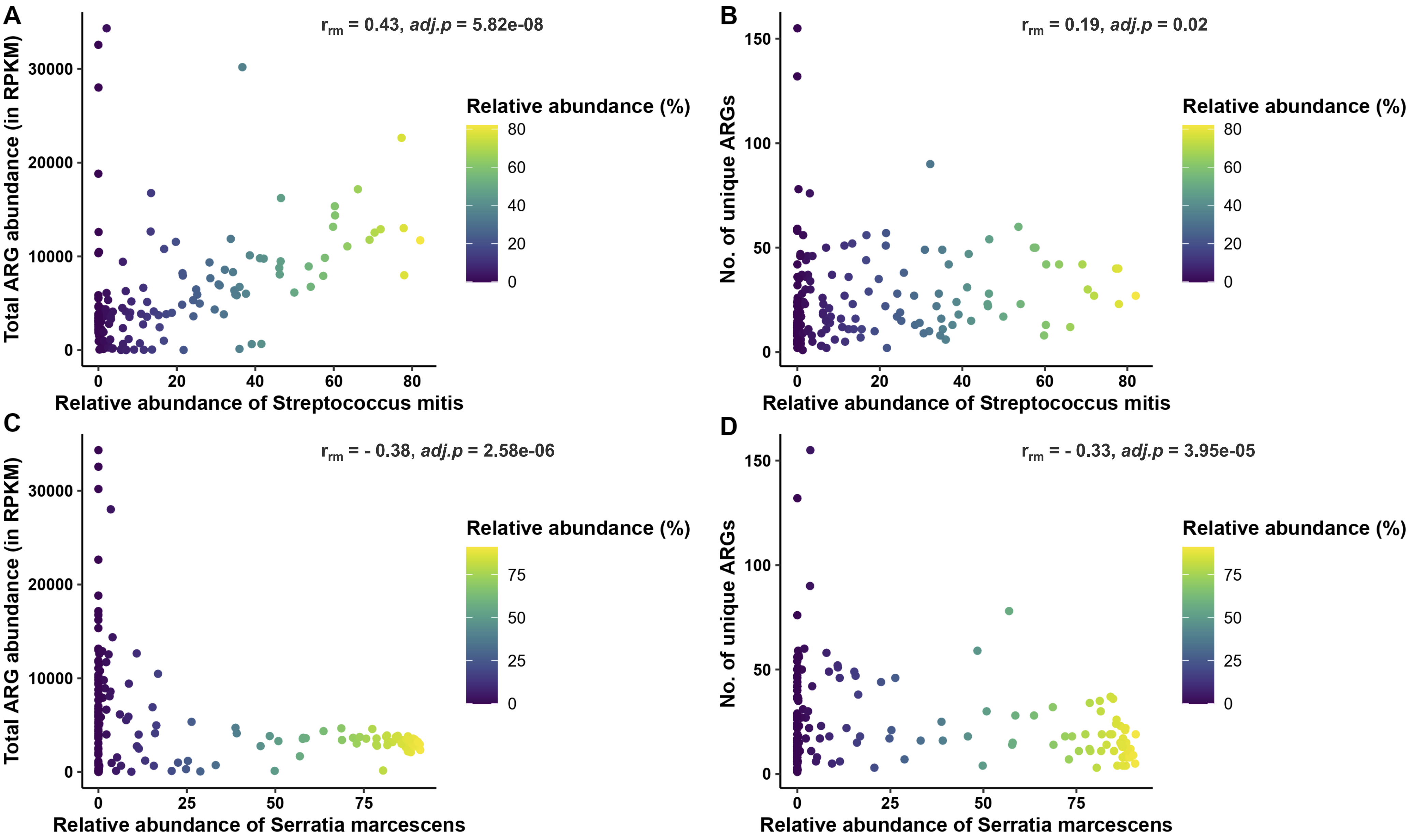

**Supplementary Fig. 9**

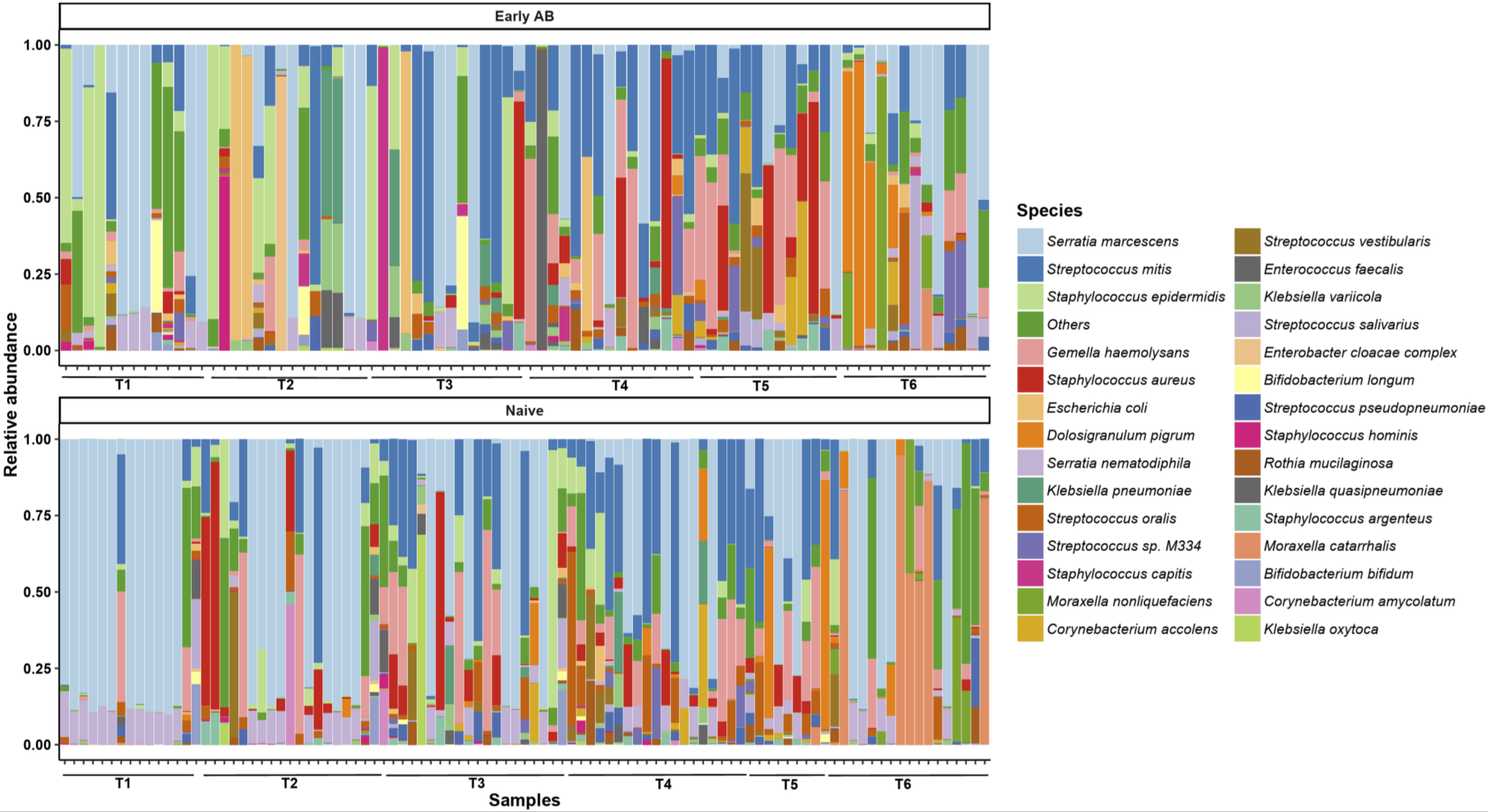
